# Sero-Prevalence and Risk Factor of *Peste Des Petits Ruminants* in Sheep and Goats of Ethiopia: Systematic Review and Meta-Analysis

**DOI:** 10.1101/2021.04.23.441083

**Authors:** Liuel Yizengaw, Wassie Molla, Wudu Temesgen

## Abstract

**Background:** Peste des petits ruminant (*PPR*) is the most common prevalent viral disease of sheep and goats that impacts productivity and international animal trade in the world and also in Ethiopia. Despite the huge economic consequences related to *PPR*, little is known about the sero-prevalence of this disease at the country levels. The objective of this systematic review and meta-analysis was to estimate a single-group summary for sero-prevalence of *PPR* disease in small ruminants of Ethiopia and assess the potential risk factor to contribute the sero-prevalence estimate.

**Methodology:** Article on *PPR* in sheep and goats were searched in PubMed, Web of Science, Google Scholar, reference lists and African online source of articles that had been conducted between 1994 to 2020 and using inclusion and exclusion criteria with restricted to those studies published in English language.

**Results:** A total of 13 published papers containing 46 district level studies were included for analyses. The single-group summary of *PPR* disease sero-prevalence in small ruminant was estimated to be 27.71% (95 % CI: 21.46 - 33.96). Overall, the estimated pooled sero-prevalence at country level in sheep was 33.56% (95% CI: 18.72–48.41) and in goats 25.14% (95% CI: 15.68–34.59). Significant heterogeneity (*I*^2^ > 80%) was noted in all pooled estimates. The visual inspection of the funnel plot demonstrated the presence of possible publication bias which could be associated with the small number of studies and longtime interval.

**Conclusions:** This quantitative review showed that the pooled sero-prevalence to be high and regional prevalence estimates of *PPR* presented here will be useful in raising awareness and advocating the Governments to engage in initiatives *PPR* control and prevention.

## Introductions

Animal rearing is an integral part of agricultural production. A total of 2.1 billion heads of small ruminants are found worldwide and they are the primary livestock resource for many poor rural families around the globe, including subsistence farmers and landless villagers as well as pastoralists [1]. Ethiopia has one of the largest livestock inventories in Africa providing support for the livelihoods of an estimated 80% of the rural poor society. The livestock sub-sector contributes about 45% of agricultural GDP, 19-20% of national GDP, and 19-20% of total exports. In Ethiopia, a total small ruminant population estimated to be about 64.04 millions of which 31.30 million of sheep and 32.74 millions of goats [2].

Though the populations of small ruminants are huge in the country; their production and productivity contribution are less. Among the limiting factor for production and productivity, infectious disease is the main constraint particularly in the developing nation like Ethiopia. Among the different infectious diseases, *Peste des petits ruminants* (PPR) is the major problem in small ruminant productivity and production. *Peste des Petits Ruminants* (PPR) is a rinderpest-like disease of goats and sheep having many common names, such as ovine rinderpest, goat plague and plague of small ruminants or Kata [3].

The disease was first described in 1942 by Gargadennec and Lalanne in the Ivory Coast, West Africa [4]. They identified the disease that was similar to but different from rinderpest in small ruminants which was not transmissible to cattle. Since then, the disease has spread far beyond its origin in Western Africa. In the past 78 years, its dissemination has been exponential and *PPR* is now present in over 70 countries across Asia, Africa, Near and Middle East, having reached Europe in 2016 (Georgia). The disease has devastating consequences on families, communities and countries [5]. The disease was first suspected in Ethiopia in 1977 following clinical observations consistent with infection with *PPR* and later the causative agent of the disease was confirmed in goats 1991. The first published account of *PPR* in Ethiopia is from 1994 and described an outbreak in goats in the capital city, Addis Ababa [6].

The *peste des petits ruminants virus* (*PPRV*) which cause *PPR is* the prototype member of the *Morbillivirus* genus in the family *Paramyxoviridae* and the order *Mononegavirales* [7]. *PPR* is characterized by fever, anorexia, necrotic stomatitis, diarrhea, mucopurulent nasal and ocular discharges, enteritis and pneumonia and occasionally sudden death [3]. It is a highly contagious animal disease affecting domestic and wild small ruminants. Once newly introduced, the virus can infect up to 90 percent of the small ruminant animal in the area, and the disease kills anywhere up to 70 percent of infected animals. It is categorized as notifiable trasboundary disease by the World Animal Health Organization (OIE) due to its potential for rapid spread and associated restrictions on the international trade of animals and animal products [8].

*PPR* is an economically significant widespread and highly contagious viral disease of small ruminants’ species. The disease spreads quickly in susceptible ruminant species, and the highest number of outbreaks occurs in sheep and goats. Cattle, camels and several wild ruminants have been infected occasionally; however, there is currently no evidence to show that the disease is maintained in these populations without concurrent infection in sheep or goats [9].

*PPR* infected and at risk countries are home to approximately 1.7 billion heads and around 80 % of the global population of sheep and goats. PPR causes annual economic losses of up to USD 2.1 billion; 30 million of animals affected every year globally; 70 countries are infected and 60% of them are from Africa. Looking beyond this figure, 5.4 billion people live in *PPR* infected area and 330 million families are at risk of losing their livelihoods, food security, and employment opportunities. Moreover, small ruminants and their products are internationally traded commodities, particularly in Africa and the Middle East; PPR considerably affects export earnings and creates supply shortages. The inability of families, communities, and institutions to anticipate, absorb, or recover from *PPR* can compromise national and regional development efforts, and turn back the clock on decades of progress [5]. The disease is currently considered as one of the main trans-boundary and notifiable disease that constitutes an emerging or re-emerging threat in many countries of the world. Considering the disease impact, March 2015, *PPR* was targeted as a high priority disease for progressive control by the World Organization for Animal Health (OIE) and the Food and Agriculture Organization (FAO) to eradicate the disease at 2030.

A comprehensive pooled sero-prevalence estimate of *PPR* has not been reported in the country, but a several individual study with wide range of prevalence estimates in sheep and goats have been reported in various regions and were very heterogeneous across regions with sero-prevalence estimates ranging from 1.7% to 85.12% [10, 11]. Reasons for the inconsistent sero-prevalence estimates of *PPR* could include an epidemic(outbreak) situation of *PPRV* in a particular geographic area, differences in methods used for identifying the disease, origin of samples, sampling strategy, and year of study, study duration and species of animal studied. An overview of knowledge on the regional sero-prevalence of *PPR* in sheep and goats will offer a better understanding of the distribution of the disease and its impacts on animal production, and will be useful in disease control.

According to Glass [12] and Dohoo *et al*., [13] systematic review and meta-analysis is a statistical technique for combining the results from several similar studies in qualitative and quantitative way respectively. The results of multiple studies that answer similar research questions are often available in the literature. It is natural to want to compare their results and, if sensible, provide one unified conclusion. This is precisely the goal of the meta-analysis, which provides a single estimate of the effect of interest computed as the weighted average of the study-specific effect estimates. When these estimates vary substantially between the studies, meta-analysis may be used to investigate various causes for this variation. Meta-analysis has been developed for summarizing the scientific evidence from the literature. This review aims to use a systematic review and meta-analysis approach to estimate the overall pooled sero-prevalence of PPR in sheep and goats from published report, and to evaluate the potential risk factors that contribute to the variability in the sero-prevalence and distribution between and within studies.

## Methodology

### Data source and literature search strategy

An optimized systemic search strategy was used to identify all published studies related to the seroprevalence of *PPR* with potential risk factors on sheep and goats in Ethiopia. Published works were searched in four electronic web search engines: PubMed, Web of Science, Reference list database, Google Scholar and African online source for articles, published between 1994 and 2020. The key words that used for searching in electronic databases were; (((Prevalence OR Incidence OR Frequency OR Detection OR Occurrence OR Identification OR Isolation OR Characterization OR Investigation OR Survey) AND (PPR OR Peste des petits ruminants OR Goat plague OR Kata OR ovine rinderpest OR Caprine rinderpest) AND (Goat OR Doe OR Buck OR Caprine OR Ovine OR Sheep OR Ram OR Ewe OR Small ruminant AND Ethiopia OR Region OR State))) in Ethiopian small ruminant. Search field option was selected as all fields. A restriction was placed on the language of publication is English.

From publications retrieve using these key words, those about the apparent sero-prevalence of *PPR* in Ethiopia were identified. First, titles and abstracts were assessed, and respective studies were examined in detail. The inclusion criteria for retrieved articles were being published in reputable journals, written in English language, cross sectional type of study design and conducted in Ethiopia, availability of the sample size with prevalence and test method, published since 1994.

### Inclusion and exclusion criteria

The review was conducted according to the systematic review and Meta-analysis guidelines Preferred Reporting Items for Systematic Reviews and Meta Analyses (PRISMA) which was developed by Moher *et al*. [14]. The guidelines were organized in different sections; on primary sort of published articles based on their title, objective and abstract. At the second stage checking type the manuscript based on list of inclusion and exclusion criteria, and decision was employed. All studies with cross sectional type of study and employed both random and non-random samplings for sample selection were considered for review. Articles selection was conducted if they were going in parallel with the inclusion criteria. Outbreak reports, cohort study papers and case control study, experimental (clinical trial) study were excluded. The third and fourth guidelines were extraction of data and meta-analysis, respectively (study screening strategy and exclusion reasons are presented (figure 1).

**Figure 1:**
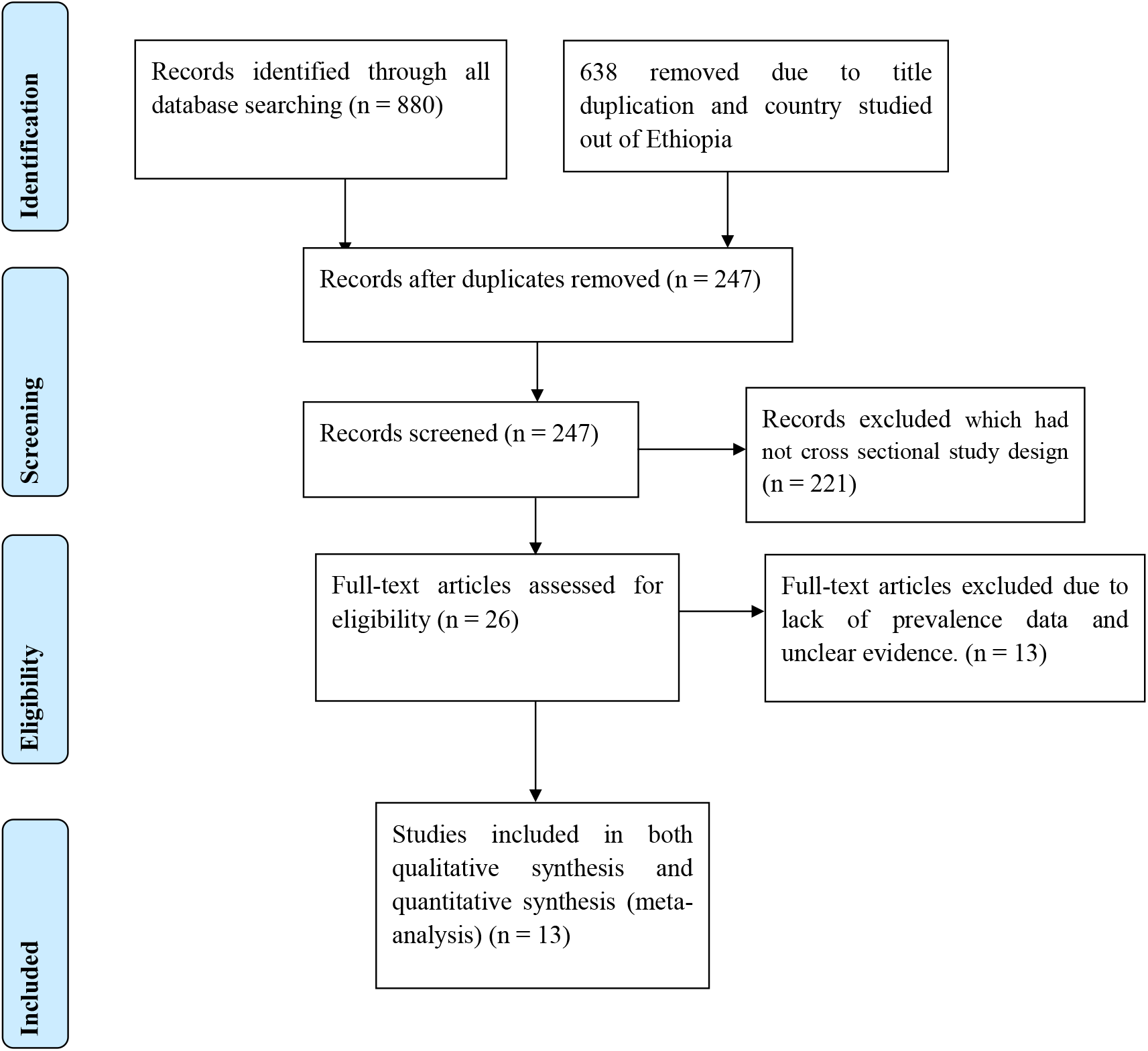
Article selection flow diagram for inclusion / exclusion of systematic review and meta-analysis process of PPR diseases.

### Data extraction

After selecting appropriate articles, variables across the study like: sampling procedure, type of design used, sex, age, district, species, administrative region, and author’s name, year of publication, type of diagnostic techniques used, sample size and apparent sero-prevalence were extracted using extraction template.

### Data analysis and management

Data analysis was conducted by using Microsoft excel 2010 and Stata software version 16 (Stata/SE 16, College Station, Texas 77845 USA. A simple summary of reports with the sero-prevalence of PPR was done by descriptive statistics. Meta-analysis of sero-prevalence data was analyzed pooled the estimates and the 95% confidence intervals. Random effects meta-analyses model for an outcome of *logit* transformed prevalence data were performed using the method of DerSimonian and Laird [15] since heterogeneity was expected. An estimate of heterogeneity between study and within the study was taken from the inverse-variance of the random-effect model [16, 13] and cross checked by calculating in excel sheet. The overall district level *logit* sero-prevalence estimate was presented by the forest plot; within the plot, the horizontal line and shaded box marks represent the confidence interval and point estimates of individual study, respectively. Subgroup analyses to determine the potential sources of heterogeneity by potential risk factors region, species, and age sex and study year. Heterogeneity between studies was evaluated through the Cochran’s Q test, *I*^2^ and τ^2^. The *I*^2^ values 0 % indicate no observed heterogeneity whereas values of 25, 50, and 75 % show low, moderate, and high degrees of heterogeneity, respectively [17]. The bias due to the use of different risk factors in the reports was assessed by using a funnel plot and subsequently Begg’s and Egger’s statistical tests. These tests were used to detect whether the bias level could statically significantly or not [18, 19]. Finally, a Meta regression model was used to quantify the source of heterogeneity between covariate variable and point sero-prevalence estimate.

## Results

### Descriptive literature search results

A total of 13 articles were eligible for the final systematic review and Meta-analysis from all screened studies. All of the eligible study have been used competitive ELISA for antibody detection. These selected eligible articles were conducted in 8 administrative regions, namely; Afar, Amhara, Benishangul-gumz, Gambella, Oromia, Somali, Southern Nation, Nationality Peoples Republic of Ethiopia (SNNRP) and Tigray from 2005 onwards. From 13 published articles and 46 study reports a total of 23961 samples of small ruminant (both sheep and goats) were subjected to disease detection at district and region level. The sample size ranges from 20 to 5992 shoat at district level and from 685 to 8321 at regional level. The sero-prevalence of the disease in the 13 articles ranges from 1.69% to 75.5%, whereas in 46 reports the sero-prevalence ranges from 1.7% to 85.1%. The highest numbers of sheep and goat sample were taken from Amhara and whereas the highest number of study reports in Oromia region as compared to others. The mean sample size from overall report was 521. Regarding to potential source of variability for the output of this systematic review and meta-analysis study in addition to region are study year which indicates that fluctuating (irregular) pattern from time to time (Figure 2), species, age and sex were used for sub group and covariate analysis. A detailed summary of the studies are presented in (Table 1).

**Figure 2:**
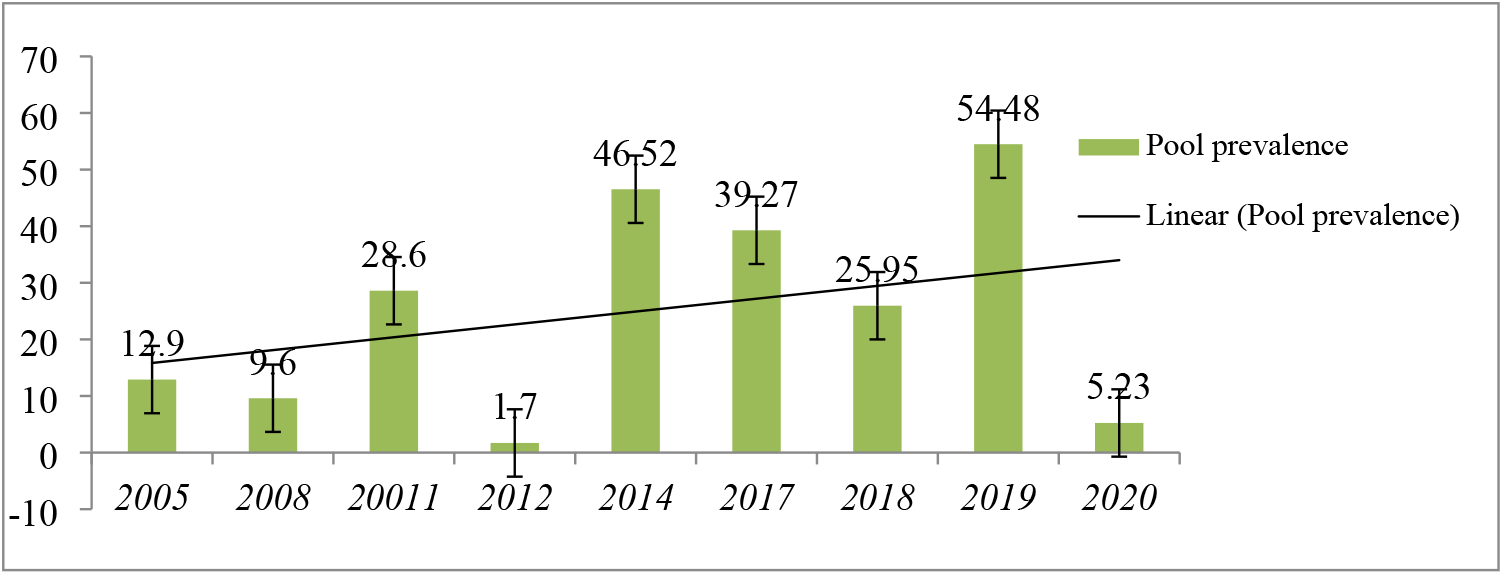
the pooled sero-prevalence trend of PPR in the meta-analysis of 46 studies between 1994-2020.

**Table 1:** Descriptive statistics of eligible studies in the final systematic review and meta-analysis.

| Authors with year | Region | District | Sample size | APP (%) | Species | SS | APP (%) | Sex | SS | APP (%) |
| --- | --- | --- | --- | --- | --- | --- | --- | --- | --- | --- |
| Abraham et al 2005 | Afar | Afar | 396 | 16 | Sheep | 835 | 23 | - | - | - |
| Abraham et al 2005 | Oromia | Oromia | 360 | 2 | Goat | 442 | 22 |  |  |  |
| Abraham et al 2005 | Oromia | Oromia | 211 | 16 |  |  |  |  |  |  |
| Abraham et al 2005 | Gambela | Gambela | 90 | 22.5 |  |  |  |  |  |  |
| Abraham et al 2005 | Somali | Somali | 220 | 11 |  |  |  |  |  |  |
| Wariet et al 2008 | Afar | - | 1653 | 15.3 | Sheep | 4211 | 8.3 | male | 1007 | 7 |
| Wariet et al 2008 | Amhara | - | 5992 | 4.6 | Goat | 4585 | 9.4 | female | 4861 | 9.4 |
| Wariet et al 2008 | Benishangul | - | 729 | 8 | Shoat | 4855 | 1.9 |  |  |  |
| Wariet et al 2008 | Oromia | - | 2290 | 1.7 |  |  |  |  |  |  |
| Wariet et al 2008 | SNNPR | - | 1622 | 1.8 |  |  |  |  |  |  |
| Wariet et al 2008 | Somali | - | 465 | 21.3 |  |  |  |  |  |  |
| Wariet et al 2008 | Tigray | - | 900 | 15.3 |  |  |  |  |  |  |
| Megersa et al 2011 | Afar | Adaar | 384 | 38.3 | Sheep | 251 | 29.5 | male | 154 | 22.1 |
| Megersa et al 2011 | Gambela | Abobo | 210 | 26.7 | Goat | 912 | 31.3 | female | 1009 | 30.1 |
| Megersa et al 2011 | Gambela | Gambela | 222 | 35.1 |  |  |  |  |  |  |
| Megersa et al 2011 | Gambela | Lare | 190 | 28.9 |  |  |  |  |  |  |
| Megersa et al 2011 | Gambela | Itang | 157 | 14.6 |  |  |  |  |  |  |
| Faris et al 2012 | Afar | Awash–<br>Fentale | 1239 | 1.7 | Sheep | 360 | 0.3 | male | 85 | 1.2 |
|  |  |  |  |  | Goat | 879 | 2.3 | female | 1154 | 1.73 |
| Afera et al 2014 | Tigray | Kukfto | 48 | 45.8 | Goat | 245 | 46.53 | male | 64 | 43.75 |
| Afera et al 2014 | Tigray | Adigudem | 48 | 50 |  |  |  | female | 181 | 47.5 |
| Afera et al 2014 | Tigray | Chercher | 48 | 50 |  |  |  |  |  |  |
| Afera et al 2014 | Tigray | Maychew | 101 | 43.6 |  |  |  |  |  |  |
| Gari et al 2017 | Oromia | Dodota | 164 | 54.9 | Sheep | 293 | 50.85 | female | 555 | 50.09 |
| Gari et al 2017 | Oromia | Adami Tullu | 353 | 45.90 | Goat | 407 | 46.68 | male | 145 | 42.07 |
| Gari et al 2017 | Oromia | Dugda | 183 | 47.54 |  |  |  |  |  |  |
| Michael et al 2017 | Oromia | Amede | 80 | 30 | Sheep | 179 | 27.9 | male | 173 | 32.37 |
| Michael et al 2017 | Oromia | Bokushenen | 128 | 35.6 | Goat | 205 | 32.1 | female | 211 | 28.44 |
| Michael et al 2017 | Oromia | Sefera | 97 | 29.6 |  |  |  |  |  |  |
| Michael et al 2017 | Oromia | Soloke | 79 | 40 |  |  |  |  |  |  |
| Gizaw et al 2018 | Afar | Adaar | 124 | 41.1 | Sheep | 94 | 41.5 | male | 60 | 45 |
| Gizaw et al 2018 | Afar | Mile | 105 | 39 | Goat | 135 | 39.3 | female | 169 | 38.5 |
| Fentie et al 2018 | Amhara | North gondar | 154 | 13.64 | Sheep | 329 | 14.9 | male | 201 | 13 |
| Fentie et al 2018 | Amhara | South gondar | 151 | 15.89 | Goat | 343 | 21.6 | female | 471 | 20.6 |
| Fentie et al 2018 | Amhara | West gojjam | 151 | 6.62 |  |  |  |  |  |  |
| Fentie et al 2018 | Amhara | East gojjam | 113 | 7.93 |  |  |  |  |  |  |
| Fentie et al 2018 | Amhara | Awi zone | 103 | 55.34 |  |  |  |  |  |  |
| Mebrahtu et al 2018 |  |  | 894 | 30.87 | Sheep | 382 | 16.2 | male | 172 | 18.6 |
|  | SNNPR | South Omo |  |  | Goat | 512 | 41.8 | female | 722 | 33.8 |
| Yalew et al 2019 | Benishangul | Bambasi | 20 | 75 | Sheep | 18 | 83.3 | male | 52 | 78.9 |
| Yalew et al 2019 | Benishangul | Homosha | 55 | 69.1 | Goat | 303 | 75.3 | female | 269 | 75.1 |
| Yalew et al 2019 | Benishangul | Sherkole | 125 | 69.6 |  |  |  |  |  |  |
| Yalew et al 2019 | Benishangul gu | Assosa | 121 | 85.12 |  |  |  |  |  |  |
| Agga et al 2019 | Amhara | N/shewa | 1657 | 2.4 | Sheep | 1055 | 11 | - | - | - |
| Agga et al 2019 | Afar | Zone three | 723 | 28.4 | Goat | 1325 | 9.6 | - |  |  |
| Gelana et al 2020 | Oromia | Horro | 152 | 3.29 | Sheep | 387 | 6.98 | male | 168 | 10.2 |
| Gelana et al 2020 | Oromia | Jimma rare | 207 | 5.31 | Goat | 419 | 4.53 | female | 638 | 11.87 |
| Gelana et al 2020 | Oromia | Jimma geneti | 447 | 6.71 |  |  |  |  |  |  |
| **Over all** |  |  | **23961** | **27.71** |  |  |  |  |  |  |

APP= apparent prevalence; SNNPR = South Nation Nationality and People of Ethiopia, SS= sample size

### Pooled and subgroup point prevalence estimate

Due to the expected variation between studies, random-effects meta-analyses were employed using the total sample size and number of positives (effect size and standard error of the effect size and also logit transformed sero-prevalence). An overall pooled prevalence of the disease was estimated to be 27.71% (21.46 - 33.96% of 95% CI) and from each reported prevalence at district level estimated ranged from 1.7% to 85.12%. Subgroup analyses were done for region (Afar, Amhara, Benishangul-gumz, Gambella, Oromia, Somali, Southern Nation, Nationality Peoples Republic and Tigray), study year (2005, 2008, 2011, 2012, 2014, 2017, 2018, 2019 and 2020), species (sheep, goat and shoat) and sex (female and male)) age (young, adult and old) of animal used (Table 2). From the subgroup Meta analyses, the estimated pooled prevalence were 25.28%, 14.82%, 60.97%, 25.55, 23.42%, 16.29%, 16.19% and 39.9% in Afar, Amhara, Benishangul-gumz, Gambella, Oromia, Somali, Southern Nation, Nationality Peoples Republic and Tigray respectively. The species, sex age classification and study year along with I^2^, Q and P value are presented in detail (in Table 2) and their sub group pooled sero-prevalence forest plot analysis result are present in figure (3, 4, 5 and 6).

**Figure 3:**
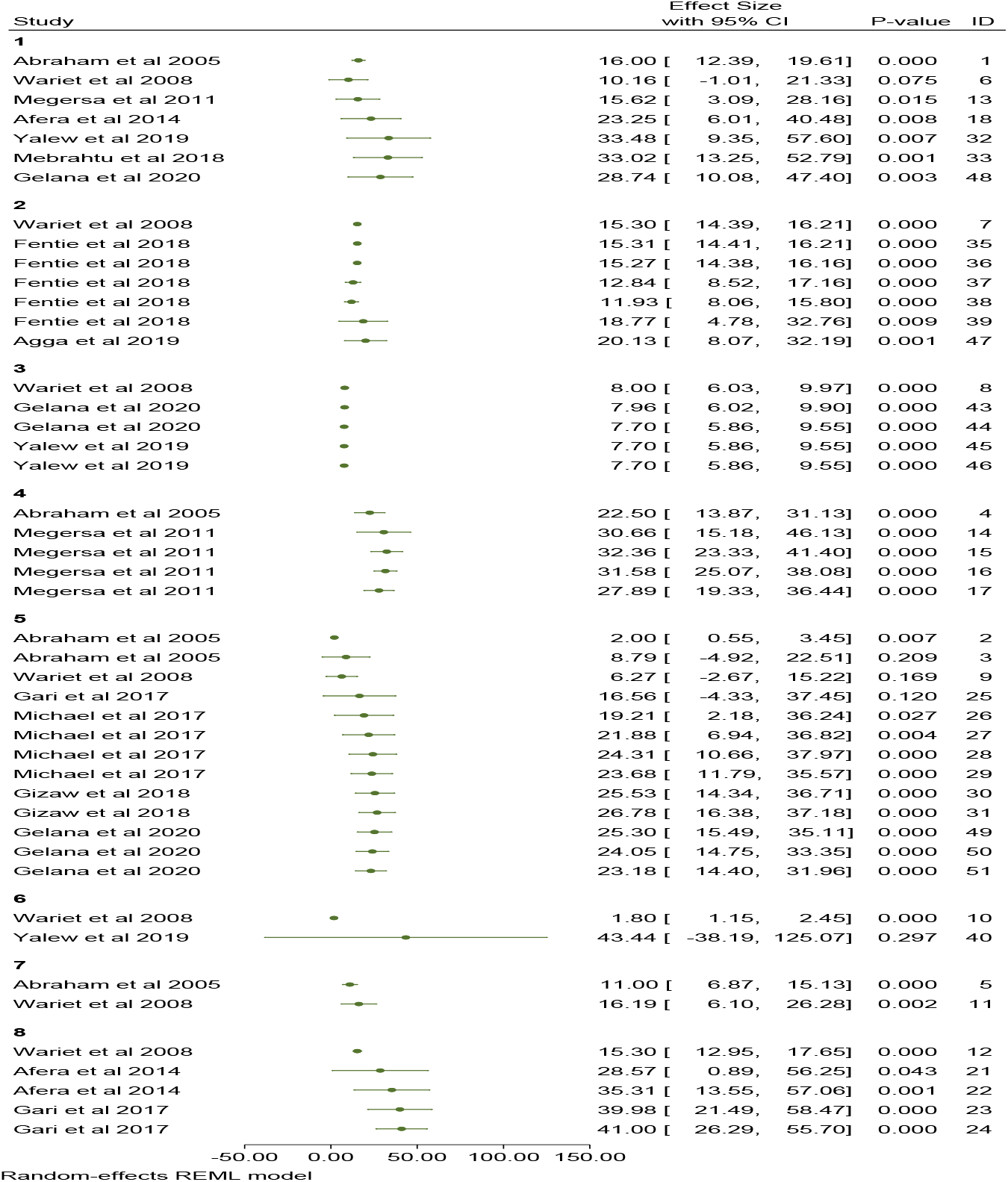
Forest plot cumulative meta-analysis by region on PPR pooled sero-prevalence estimates in Ethiopia.

**Figure 4:**
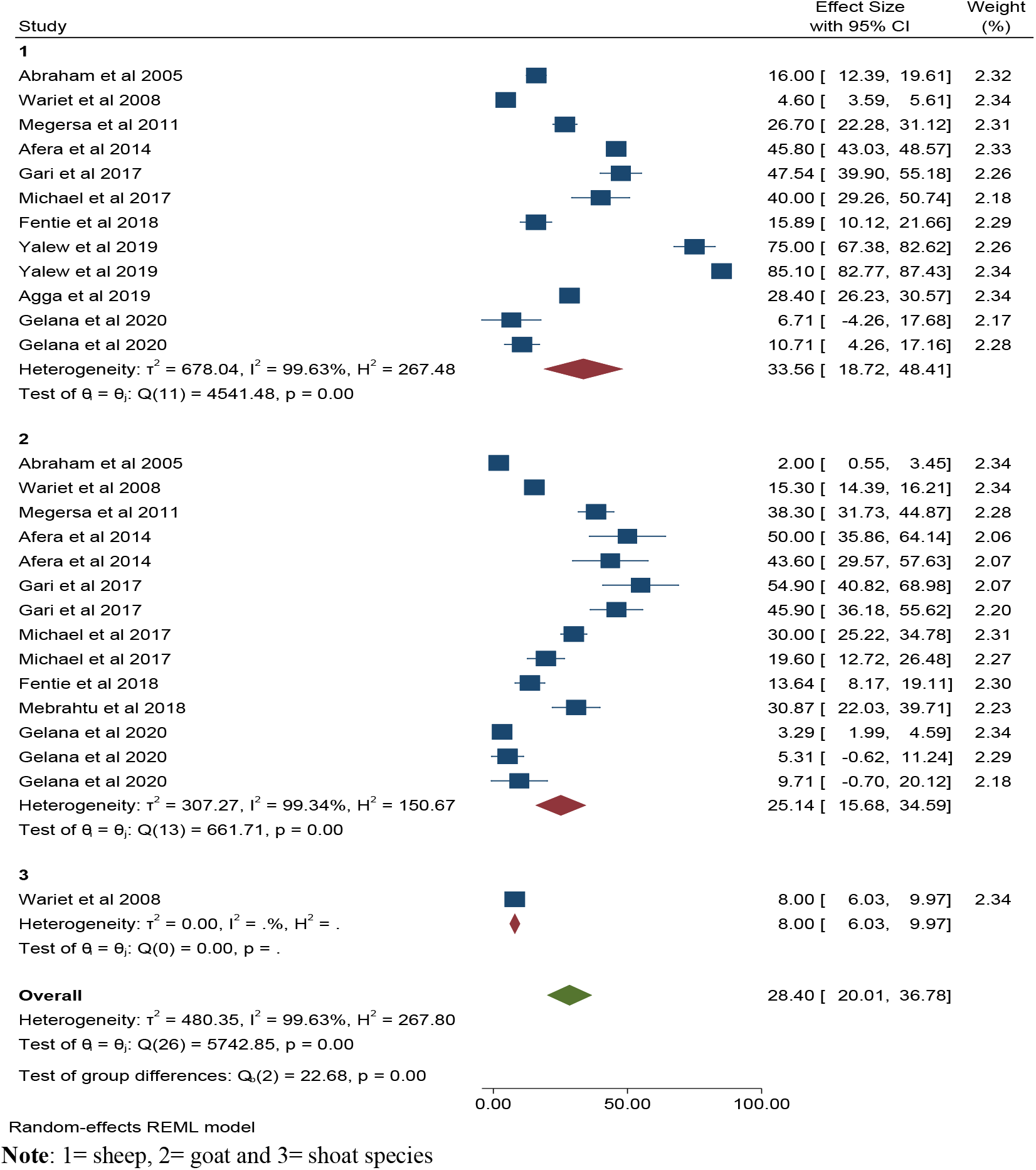
Forest plot sub group analysis by species on PPR pooled sero-prevalence estimates in Ethiopia.

**Figure 5:**
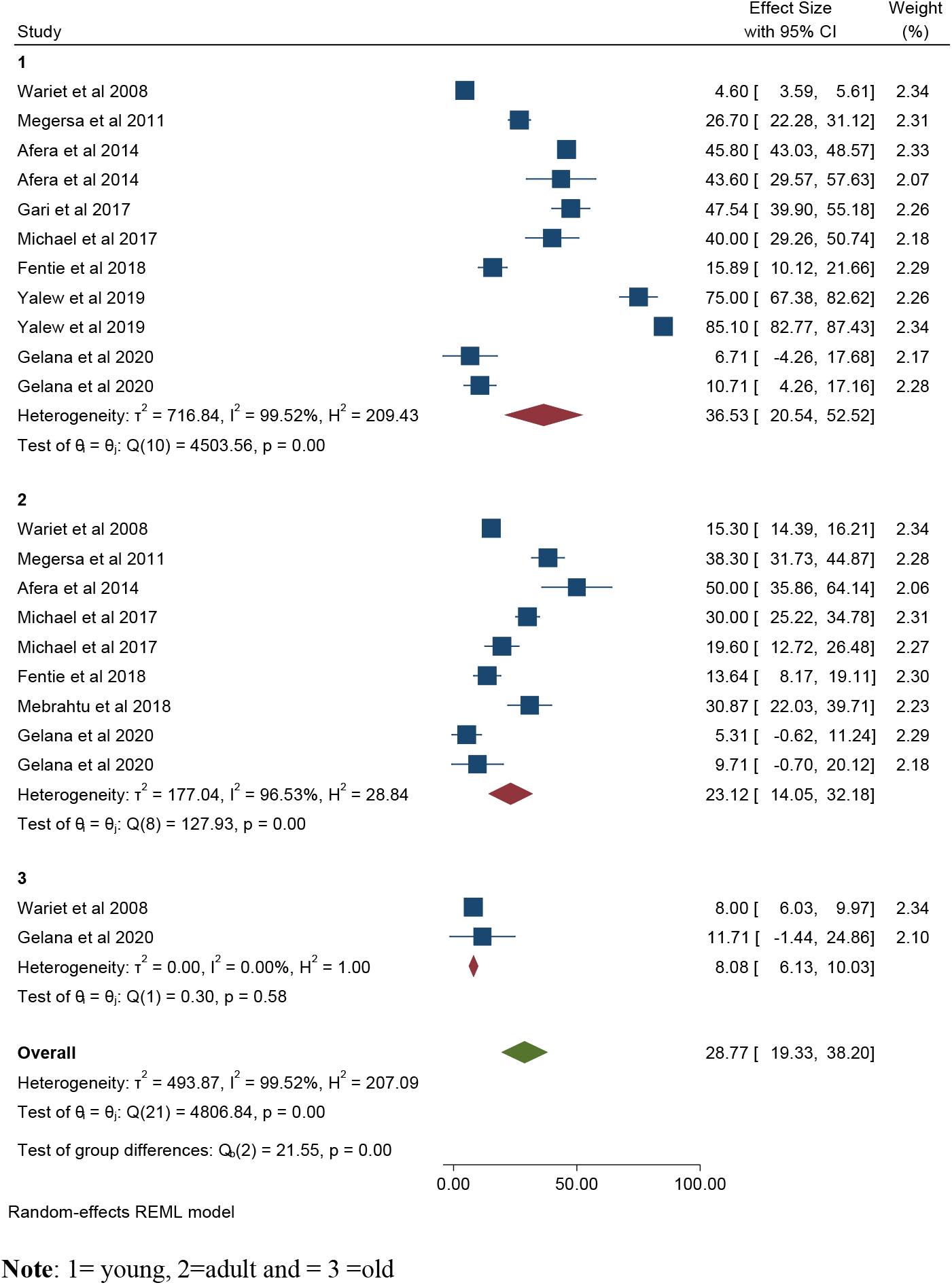
Forest plot sub group analysis by age category on PPR pooled prevalence estimates in Ethiopia.

**Figure 6:**
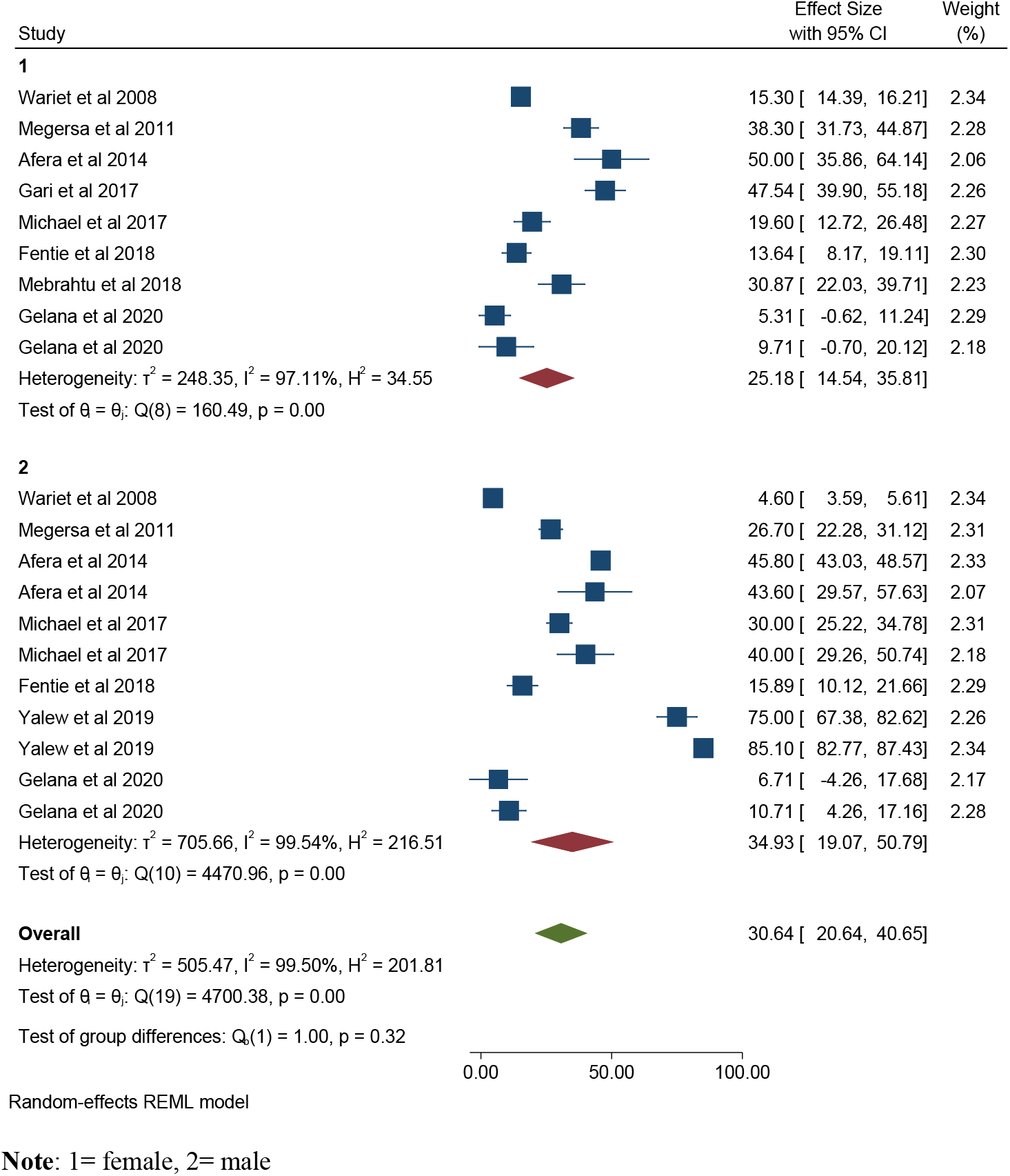
Forest plot sub group analysis by sex on PPR pooled sero-prevalence estimates in Ethiopia.

**Table 2:** Subgroup pooled sero-prevalence estimate of PPR with I^2^, Q and P value.

| Variable | Category | Sero-prevalence % (95 CI) | $I^2$ | $Q$ | DF | P-value |
| --- | --- | --- | --- | --- | --- | --- |
| Regions | Afar | 25.28(14.12-36.46) | 99.26% | 1210.45 | 6 | <0.001 |
|  | Amhara | 14.82(1.75-27.89) | 99.81% | 220.81 | 6 | <0.001 |
|  | Benishangul-gumz | 60.97(33.56- 88.38) | 98.77% | 0.75 | 4 | <0.001 |
|  | Gambella | 25.55(18.64-32.46) | 82.61% | 38.41 | 4 | <0.001 |
|  | Oromia | 23.42(12.78- 34.06) | 99.54% | 567.27 | 12 | <0.001 |
|  | SNNPR | 16.29(-12.19-44.79) | 99.70% | 4543.21 | 1 | <0.001 |
|  | Somali | 16.19(6.09-26.28) | 92.41% | 13.17 | 1 | <0.001 |
|  | Tigray | 39.9 (25.87-53.94) | 90.97% | 94.08 | 4 | <0.001 |
|  | **Overall** | 27.706(21.45-33.95) | 99.81% | 3720.06 | 45 | 0.000 |
| Study year | 2005 | 12.9 (6.3-19.59) | 93.87% | 93.94 | 4 | <0.001 |
|  | 2008 | 9.6 (3.9-15.2) | 99.63% | 468.32 | 6 | <0.001 |
|  | 2011 | 28.6 (20.57- 36.6) | 89.96% | 44.36 | 4 | <0.001 |
|  | 2012 | 1.70(0.980-2.42) | - | 0 | 0 |  |
|  | 2014 | 46.52(40.28 52.76) | 0% | 0.83 | 3 | 0.843 |
|  | 2017 | 39.27(30.38-48.15) | 89.03% | 53.81 | 6 | <0.001 |
|  | 2018 | 25.95(13.68-38.21) | 97.62% | 211.85 | 7 | <0.001 |
|  | 2019 | 54.48(28.33-80.64) | 99.55% | 1257.53 | 5 | <0.001 |
|  | 2020 | 5.23(3.18-7.27) | 41.28% | 3.35 | 2 | 0.187 |
|  | Overall | 27.706(21.45-33.95) | 99.81% | 3720.06 | 45 | <0.001 |
| Species | Sheep | 33.56(18.72-48.41) | 99.34% | 4541.48 | 12 | <0.001 |
|  | Goat | 25.14(15.68-34.59) | 99.63% | 661.71 | 13 | <0.001 |
|  | Shoat | 8.00(6.03-9.97) | - | 0.00 | 0 | <0.001 |
|  | Overall | 28.39(20.01-36.79) | 99.8% | 5742.85 | 26 | <0.001 |
| Age | Old | 8.08(6.13-10.03) | 0% | 0.30 | 1 | 0.585 |
|  | Adult | 23.112(14.05-32.18) | 96.53% | 127.93 | 8 | <0.001 |
|  | Young | 36.53(20.54-52.52) | 99.52% | 4503.56 | 10 | <0.001 |
|  | Overall | 28.76(19.33-38.20) | 99.52% | 4806.84 | 21 | <0.001 |
| Sex | Female | 25.17(14.54-35.81) | 97.115 | 160.49 | 8 | <0.001 |
|  | Male | 38.56(20.25-56.85) | 99.635 | 4451.03 | 9 | <0.001 |
|  | Over all | 30.64(20.64-40.65) | 99.50% | 4700.38 | 19 | <0.001 |
|  | **Overall** | **27.7(21.46 - 33.96)** | **99.81%** | **3720.06** | **45** | **<0.001** |

**Note:** 1, 2, 3, 4, 5, 6, 7, and 8, represent Afar, Amhara, Benishangul-gumuz, Gambella, Oromia, Somali, Southern Nation, Nationality Peoples Republic of Ethiopia and Tigray region of the study reported respectively.
Q=Heterogeneity chi-square, $I^2$ = effect size residual variation due to heterogeneity, DF= degree of freedom.

### Meta-analysis result of *PPR* diseases by authors

The effect size meta-analysis result of pooled sero-prevalence at district level was 27.71 (95% CI: 21.46- 33.96). The analysis results indicated that between-study variability was high (τ^2^ =454.02; heterogeneity I^2^ = 99.81% with Heterogeneity chi-square (Q) = 3720.06, with degree of freedom 45 and P value of 0.000 (Figure 7). Studies weighted almost equal with weights on individual studies ranging from 1.86% to 2.24% due to high heterogeneity between studies presents the forest plot derived from the meta-analysis with the effect size and corresponding weight for each risk factor. The effect size of the pooled *logit* prevalence result of random effect meta-analysis was −2.8 (95% CI: −3.33 to −2.28). The *logit sero*-prevalence estimate also indicates a moderate proportion of between study variance (*I*^2^=42.8%, τ^2^=0.65) (Figure 8). In respected to individual author study, the meta-analysis result of pooled sero-prevalence indicated that high variability with τ^2^= 453.67; I^2^ = 99.88% with Cochran’s Q statistics 2524.17, at a degree of freedom 12 with a P value of = 0.00. Individual study sero-prevalence estimates ranged from 1.69% to 75% with the overall random pooled sero-prevalence of 27.5% (95% CI: 15.88 - 39.12) (Figure 9).

**Figure 7:**
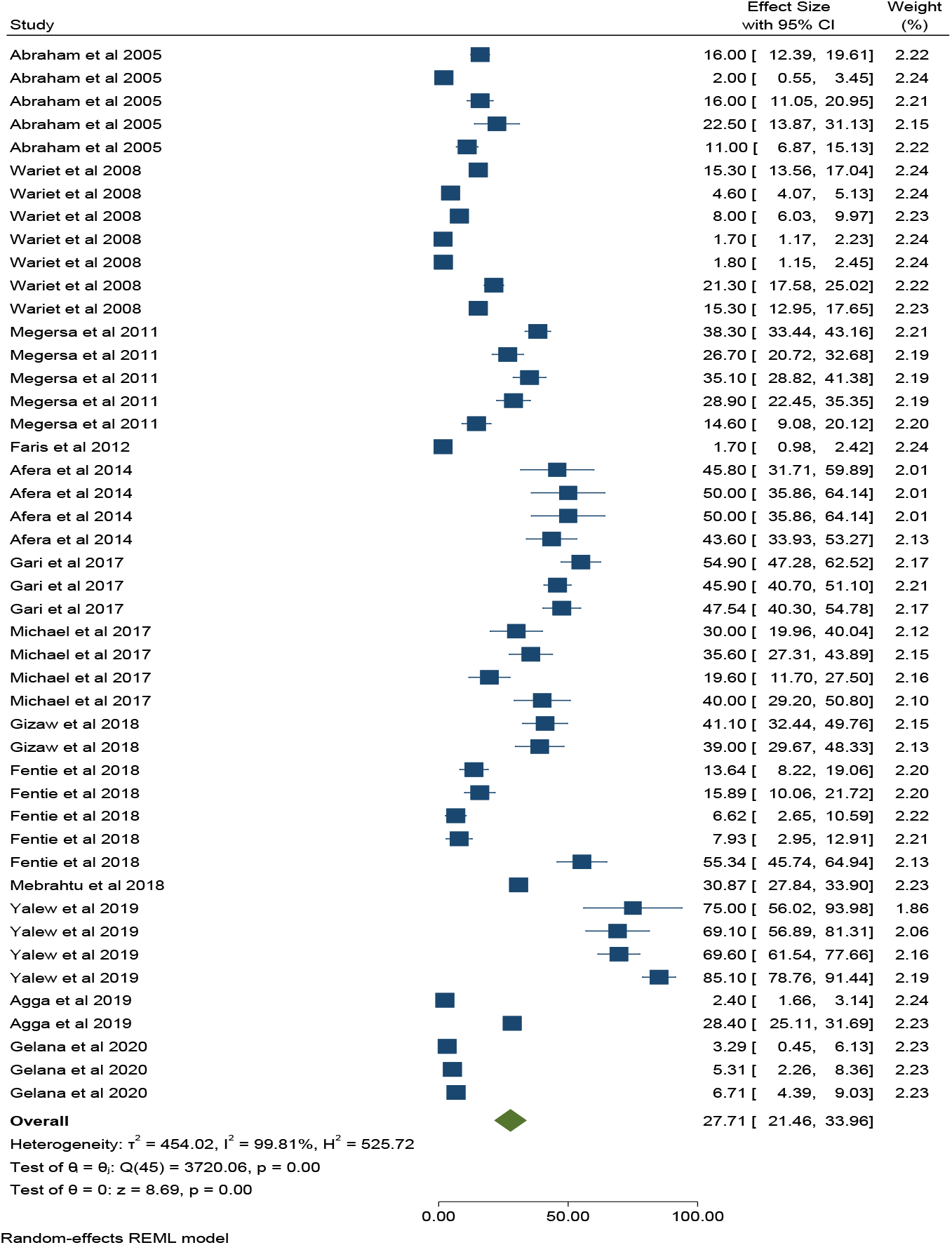
Forest plot on PPR of sheep and goats pooled sero-prevalence estimates in Ethiopia at district level (46).

**Figure 8:**
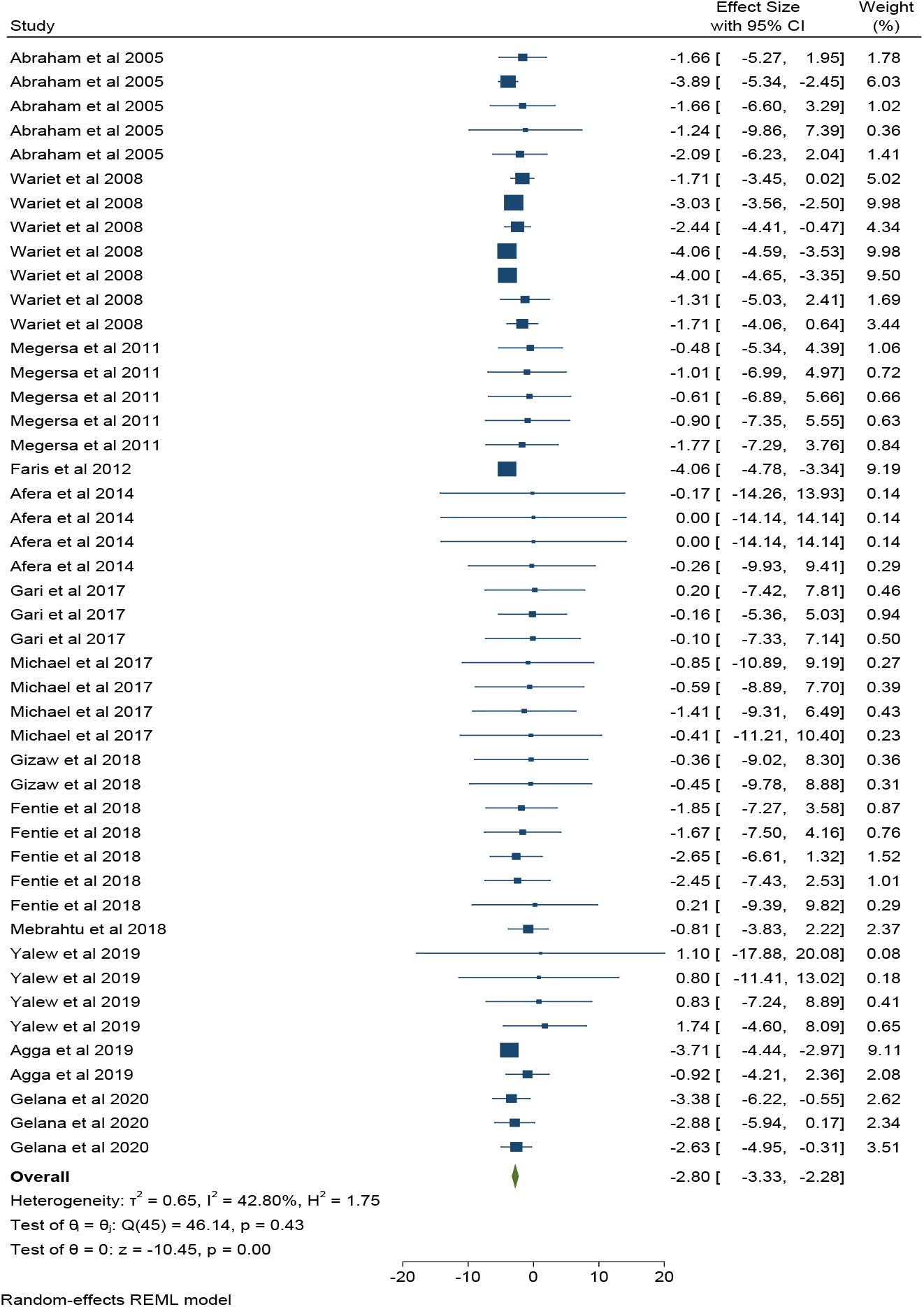
Forest plot on PPR of Shoat pooled logit-sero-prevalence estimates in Ethiopia at district level (46)

**Figure 9:**
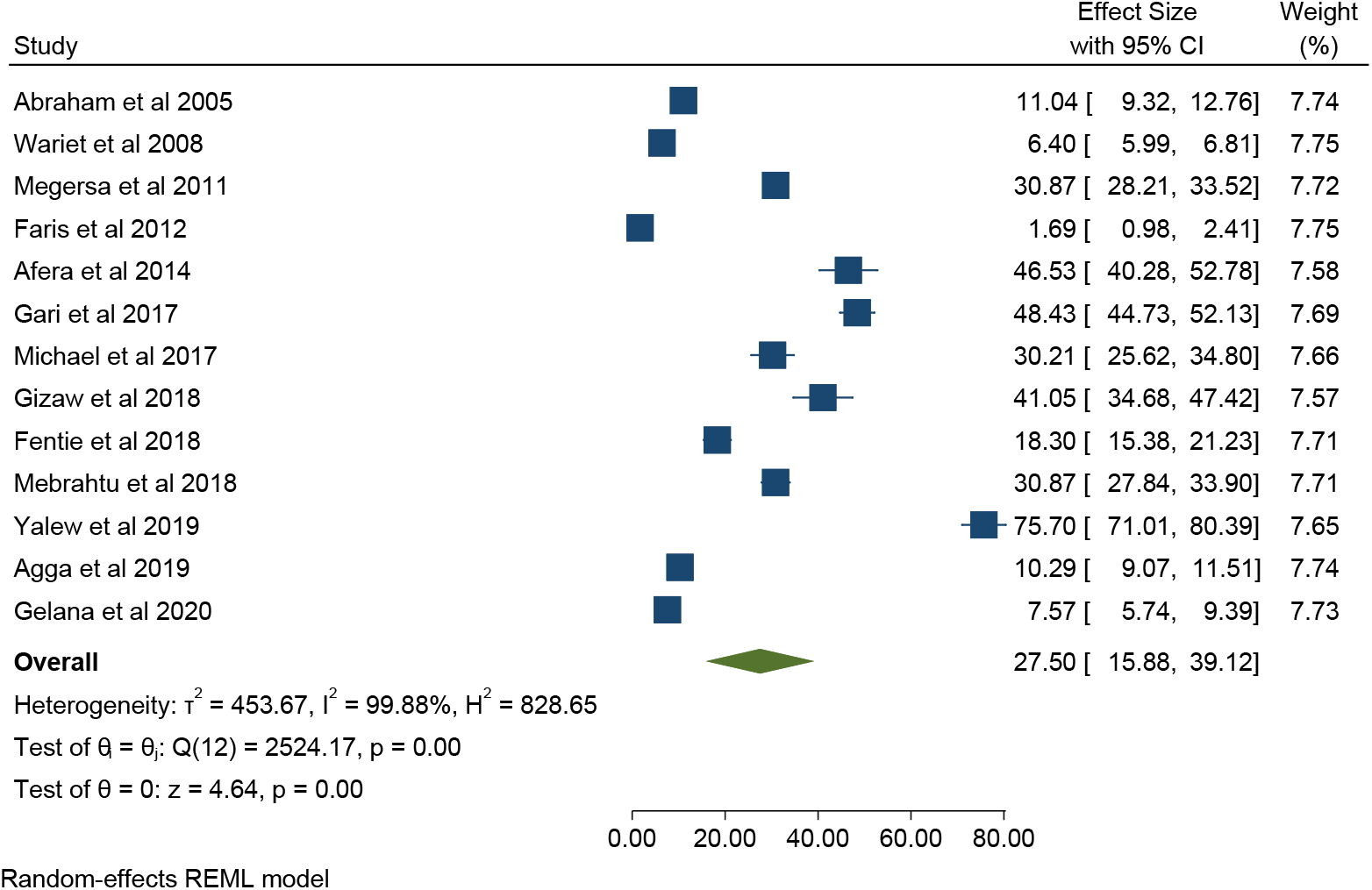
Forest plot on PPR of shoat pooled sero-prevalence estimates in Ethiopia by individual authors study.

### Univariable and multivariable meta-regression

Results with coefficients, *p* values, I^2^ and τ^2^ from multivariable and univariable meta regression are presented in Table 3. All predictor variables in both univariable and multivariable meta regression analysis were not statistically significant (p >0.05).

**Table 3:**
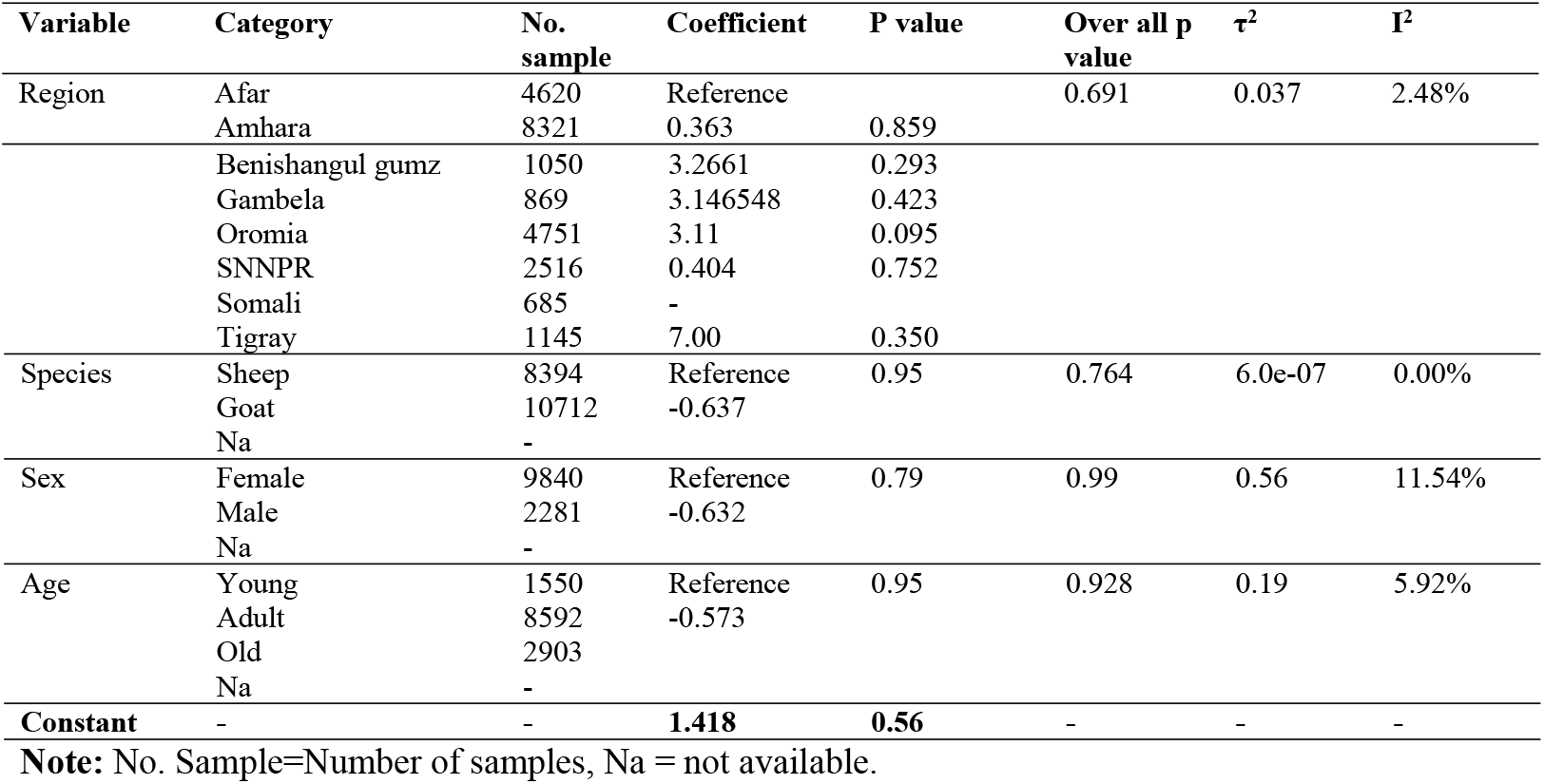
Univariable and multivariable regression coefficient on logit sero-prevalence estimate on PPR disease of small ruminant in Ethiopia (46 district level reports)

### Publication bias results

From our assessments of bias and small study effects by funnel plot observation for small-study effects. The result of effect estimates against its standard error showed that there was a publication bias with a p-value of <0.001 (Fig. 10). From Begg’s and Egger’s test statistics result there was no any study effect since the estimated bias coefficient 4.77 with standard error 0.198.

**Figure 10:**
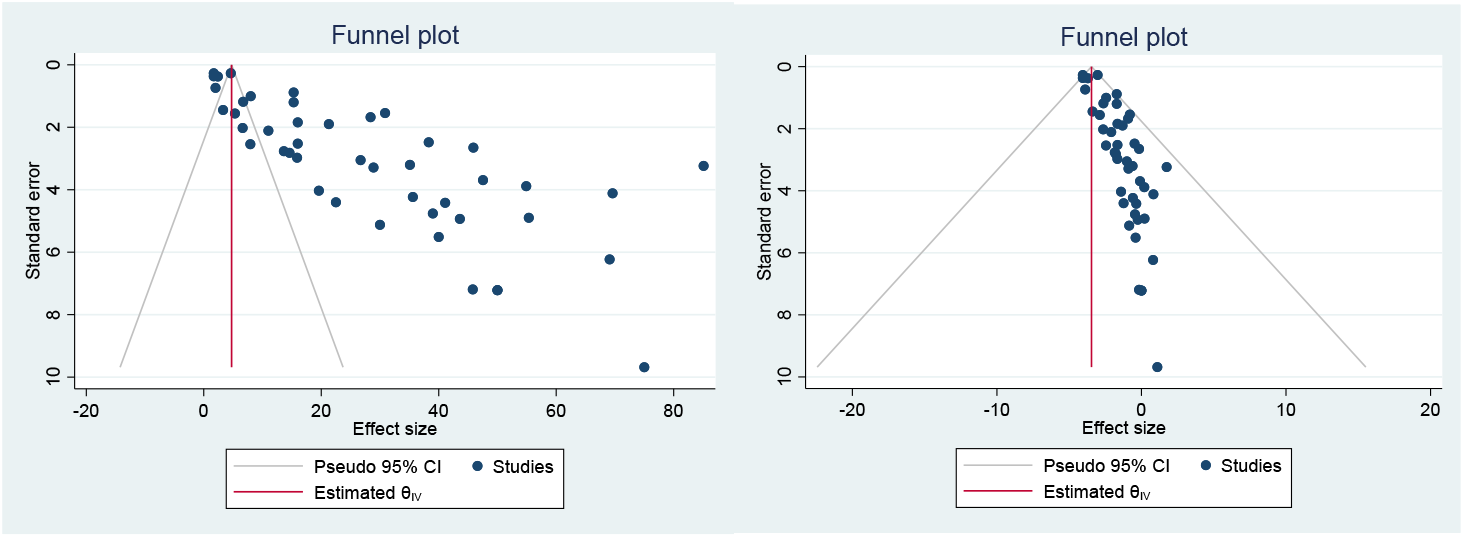
Funnel plot that assesses publication bias (left from estimated pool sero-prevalence: right side from pooled logit sero-prevalence effect size.

## Discussion

*Peste des petits ruminants* (*PPR*) continue to cause the death of millions of sheep and goats annually and are a constant threat to the livelihoods of subsistence farmers in many countries in Africa, the Middle East and Asia. In Ethiopia a number of cross sectional studies and outbreak reports are available across different agro ecology of the country in different year. But the sero-prevalence of the disease is affected by different factors like, environmental factors, the number of samples, type of strains, stage of infection and type of diagnostic techniques used. The approaches of Meta-analysis allow identifying the role of such factors, by combining results of different reports, with different designs, agro ecology and locations. Good meta-analysis outputs are relevant for the management and control of an infectious disease like *PPR* that could not be identified by individual studies alone [13]. This is the first quantitative meta-analysis on the sero-prevalence and risk factors of *PPR* in sheep and goats in Ethiopia to the best of theour knowledge. We have used 46 point prevalence reports from 13 cross sectional studies that have been undertaken between year 2005 and 2020.

Country wise the reported apparent sero-prevalence of *PPR* ranges from 1.7% to 85.12% [10, 11]. Such variability is attributed to the situation of *PPRV* in a particular geographic area, differences in methods used for identifying the disease, origin of samples, sampling strategy, and stage of infection, study duration and species of animal and the size of sample used. The Meta analysis of the effect size of the random effect model considers the existing real difference between study reports beyond chance. The value of the inverse variance square (99.81%) revealed the sero-presence of true variability and high heterogeneity. But in the Meta regression the other factor did not contribute additional heterogeneity among studies.

The pooled sero-prevalence estimate varies significantly between regions and higher in Benishangul-gumz (60.97%) and Tigray 39.9% as compared to other regions. The difference in sero-prevalence could be attributed to ecological characteristics of a specific area, such as climate, settlement pattern, sanitary and socio-economic practices. In both Benishangul-gumz and Tigray region are adjacent with the border and may be getting an access of infection from abroad country. This report in line with Ahaduzzaman [20] explained that the variation of the disease prevalence like *PPR* could be the trans boundary movement of infected animals with inadequate quarantine, the presence of hot and humid climatic conditions that favor disease epidemiology, lack of vaccination or vaccine administration monitoring which may facilitate disease spread, lack of awareness about *PPR* among backyard farmers, and limited funding for disease eradication in developing or underdeveloped countries. Moreover many studies included in this meta-analysis used serum sample or symptomatic diagnostic approaches to report *PPR* prevalence; such approaches can quickly reveal the status of a large population [10, 11] could lead to variability across region.

The estimated pooled sero-prevalence indicated that *PPR* is varies significantly between species and higher in sheep 33.56% than goats 25.14%. This report is similar with other several studies [21, 22, 23, 24, and 25] and disagreed with reports that indicated *PPR* is more prevalent in goat than sheep [10, 26, 27]. Although there are biological differences between sheep and goats, higher sero-prevalence in one species than another could be due to factors such as sampling process, richness or distribution of animal in a geographical area, management practices and strain of the virus. It is also possible that *PPRV* preferentially infects goats over sheep or vice-versa depending on the prevailing situation and the disease severity may also vary between species [28].

*PPR* was significantly higher in young animals (36.53%) than in adult (23.12%) and old (8.08%) animals based on the estimated pooled prevalence. These results agree with the findings of many studies [29, 30, 31, 32], but it is not in line with many other reports [21, 33, 34]. The higher prevalence in young animal than the old could be due to malnutrition, less developed immune system and poor husbandry practices [31, 35]. It has been reported that *PPRV* is highly immunogenic, and animals remain seropositive for a long period, particularly in an endemic area [33, 36].

The estimated pooled sero-prevalence of PPR was significantly higher in male (38.56%) than in female (25.17%) animals. This could be due to the proportion of sampled animal during the study. This report is in line with other studies report [11, 24, 29, 37]. Rony *et al* [37] reports describe that a higher prevalence in males, possibly due to a higher proportion of male animals in a flock particularly when the age of the studied animals was under two years. And the high demand of male animals for meat purpose driven them to the market and contribute to the higher infection rate than female which kept at home for breeding purpose and also due to genetic variation of the animals [38]. In contrast many other authors reports higher prevalence in female than male [10, 22, 25, 26, 27, 34, 39, 40, 41, 42, 43]. This could be due to breeding females being used for flock reproduction maintenance for a more extended period than males and higher density of females than males in flocks, or physiological differences between females and males

The report of Egger’s test statistics and inspection of the funnel plot revealed that there is an evidence for presence of publication bias. However, the source of funnel plot asymmetry could additionally be due to true heterogeneity or unable to incorporate unpublished reports and it may be small number cross sectional reports used or even to chance [19].

### Limitations

The study has some limitations; an overall analysis of the study showed a large degree of heterogeneity among studies and within subgroup analysis. The studies used in this analysis used only competitive ELISA for diagnosis which has low precision for detection of antibody as compared to PCR and others. Moreover, they luck full information on important factors like; species age, and the sex of the animal properly. The absence of unpublished data in the Meta analysis also limits the reflection on the real epidemiology of the disease in the country. And many report found in the form of outbreak and the study covers long period of time since 2005 to 2020 due to insufficient cross sectional data. Therefore the study may not necessarily reflect the real situation of the country disease status.

## 5. CONCLUSION AND RECOMMENDATIONS

This is the first systematic review and meta-analysis study made on prevalence and risk factors of *PPR* disease in sheep and goats of Ethiopia. The pooled prevalence estimate of the disease is high even though higher degree of variability was observed among studies, between regions, and associated risk factors. The disease was found to be more prevalent in sheep young and male animals thus preventing method like vaccination in both species may prevent the disease spread and enhance to achieve the eradication goal. To get clear picture of the prevalence of the disease in the country, an outbreak data should be combined. Additionally, factors that contribute to the prevalence estimate heterogeneity should be handled appropriately in any cross sectional study to accurately estimate the true extent of *PPR sero-prevalence*.

## Registration

No register name and registration number since it is scoping review

